# Chemical-state-dependent free energy profile from single-molecule trajectories of biomolecular motor: Application to processive chitinase

**DOI:** 10.1101/655878

**Authors:** Kei-ichi Okazaki, Akihiko Nakamura, Ryota Iino

## Abstract

The mechanism of biomolecular motors has been elucidated using single-molecule experiments for visualizing motor motion. However, it remains elusive that how changes in the chemical state during the catalytic cycle of motors lead to unidirectional motions. In this study, we use singlemolecule trajectories to estimate an underlying diffusion model with chemical-state-dependent free energy profile. To consider nonequilibrium trajectories driven by the chemical energy consumed by biomolecular motors, we develop a novel framework based on a hidden Markov model, wherein switching among multiple energy profiles occurs reflecting the chemical state changes in motors. The method is tested using simulation trajectories and applied to singlemolecule trajectories of processive chitinase, a linear motor that is driven by the hydrolysis energy of a single chitin chain. The chemical-state-dependent free energy profile underlying the burnt-bridge Brownian ratchet mechanism of processive chitinase is determined. The novel framework allows us to connect the chemical state changes to the unidirectional motion of biomolecular motors.

## INTRODUCTION

The single-molecule experiment is a powerful tool for studying the mechanism of biomolecular motors via motion visualization. Various single-molecule techniques have been used to elucidate the mechanisms of F_1_-ATPase^1–6^, V_1_-ATPase^7,8^, myosin^9–12^, kinesin^13–15^ and dynein^16–19^, the typical motors driven by adenosine triphosphate (ATP). Recently, single-molecule experiments have been used to identify processive cellulase and chitinase as a different type of biomolecular motor, in which the hydrolysis energy of polysaccharides is used for unidirectional motion^20–24^. These experiments have been combined with structural information and molecular dynamics simulations to elucidate how changes in the chemical state during the catalytic cycle are coupled to structural changes and mechanical motions^22,25–30^. However, experiments and simulations are not able to track the structural dynamics of active motors at atomic detail over timescale relevant for unidirectional motions. Thus, simplified mechanisms have been proposed to explain how the motors generate unidirectional motions: the power stroke and the Brownian ratchet mechanisms^31,32^. Although models based on these two mechanisms have been developed^33–41^, single-molecule experimental data have not been fully exploited to develop realistic mechanisms.

Attempts have been made to estimate physicochemical models directly from single-molecule trajectories of equilibrium processes such as conformational dynamics. Single-molecule fluorescence (or Förster) resonance energy transfer (FRET) trajectories of DNA, RNA and protein conformational dynamics have been analyzed using the hidden Markov model (HMM) or the maximum likelihood method to estimate the kinetic rates and FRET efficiencies associated with the different conformational states that produce FRET signals^42–47^. FRET trajectories have also been used to estimate Langevin dynamics^48^. Single-particle tracking trajectories of a cell-surface receptor have been analyzed by HMM to estimate multiple diffusion coefficients corresponding to different conformational states of the receptor^49^. The energy landscape is another key quantity for understanding the motion, kinetics and stability of biomolecules^50–54^ that can be estimated from single-molecule trajectories^55,56^. Along the lines of these studies, we aim to develop a method that is applicable to nonequilibrium processes, such as the single-molecule trajectories of biomolecular motors.

Biomolecular motors move unidirectionally along a linear track or rotate relative to a stator by consuming chemical energy. Thus, motor motion can be effectively described by a onedimensional (1D) model. This simplification has been employed for rotary^57,58^ and linear^37^ motors. A similar approach has been explored in the field of protein folding. Apparently complex processes can be projected onto a 1D reaction coordinate and modeled as a diffusion process over an energy barrier separating unfolded and folded states^45,59–61^. Methods have been developed for construction of a 1D diffusion model from experimental or simulation trajectories^62,63^. In particular, the method developed by Hummer and coworkers has been well tested with extensive applications to simulation trajectories^59,60,62,64^. However, it is challenging to apply the methods to biomolecular motors, because the unidirectional motor motions generally involve switching among different energy profiles depending on the motor chemical states.

In this study, we develop a method to construct a 1D diffusion model that involves switching among multiple energy profiles from the trajectories of biomolecular motors. In the novel framework, Hummer’s method is combined with HMM to estimate the chemical-state-dependent energy profile and the diffusion coefficient. We test the method using trajectories generated from a Brownian dynamics simulation in which two energy profiles switch stochastically. Then, we apply the method to single-molecule trajectories of processive chitinase to obtain the chemicalstate-dependent free energy profile triggered by the hydrolysis reaction of a single chitin chain.

## METHODS

### Likelihood function for diffusion model with multiple energy profiles

Here, we develop a framework for estimation of the diffusion model that involves switching among multiple energy profiles. The underlying concept of the framework is that among different hidden states, each with their own energy profiles and diffusion coefficients, one of these states is adopted at each snapshot of a trajectory. As an example, we consider a trajectory with segments that are either in state 1 or 2 (Fig. 1A, red or blue). In addition to estimating the energy profiles and diffusion coefficients, we must determine the state that is adopted at each snapshot of the trajectory. This type of problem can be solved by HMM^65^. Given a model, the likelihood of a trajectory *X* = (*x*_0_,*x*_1_,…,*x_N_*) is

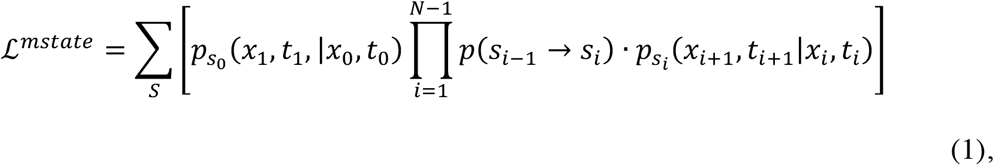

where 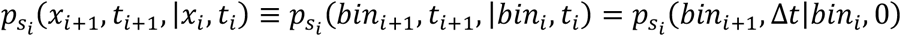 is the positional transition probability in state *s_i_*, discretized into bins with lag time Δ*t* = *t*_*i*+1_ – *t_i_*, and *p*(*s*_*i*-1_ → *s_i_*) is the switching probability from state *s*_*i*-1_ to *s_i_* in time Δ*t*. All possible paths of the hidden states *S* = (*s*_0_, *s*_1_, …,*s*_*N*-1_) are summed over. The positional transition probability *p_s_i__*. (*bin*_*i*+1_, Δ*t*|*bin_i_*, 0) was computed in the same manner as the single energy profile case developed by Hummer and coworkers^60,62,61^ (see below). Direct calculation of this likelihood is difficult because the number of the possible hidden-state paths grows exponentially with *N* (the number of snapshots in a trajectory). However, the likelihood can be calculated using the HMM forward-backward algorithm at a computational cost ~*O*(*N*) (see SI text).

**Figure 1.**
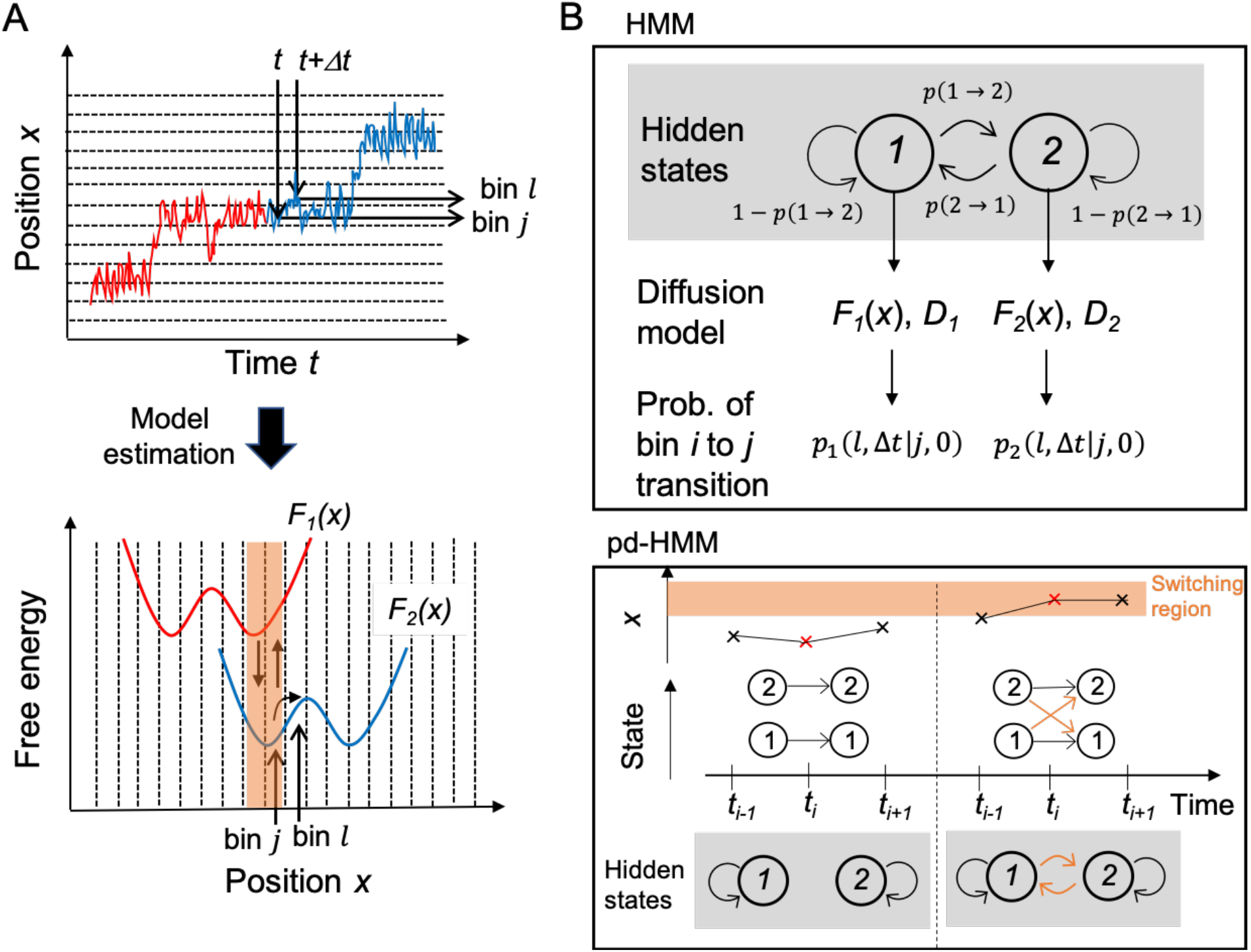
Estimation of chemical-state-dependent free energy profile. (A) Estimation of underlying free energy profiles and diffusion coefficient from one-dimensional trajectories, where red and blue segments of trajectory in top panel correspond to diffusion over red and blue energy profiles in bottom panel, respectively. Orange region represents the switching region. (B) Top panel shows HMM scheme to construct diffusion model with two energy profiles. Bottom panel shows position-dependent HMM scheme that is an extension of traditional HMM.

In the HMM context, the observed data are positional transitions between bins, the state transition probability corresponds to the switching probability *p*(*s*_*i*-1_ → *s_i_*), and the output probability of the particular transition bin *j* → bin *l* in state *s_i_* is proportional to *p_s_*(*l*,Δ*t|j*, 0) (see top panel of Fig. 1B). As the output probabilities must sum to unity, the positional transition probabilities are divided by the number of bins and used as the output probabilities. Given a model (that is, an energy profile and diffusion coefficient for each state as well as switching probabilities), the likelihood 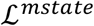 and the state probability *p_t_i__*(*s*) ≡ *p*(*s_i_* = *s*|*X*) at any point in the trajectory can be computed by the HMM forward-backward algorithm. The state probability *p_t_*(*s*) indicates which energy profile is likely to be adopted at time *t*. To deal with multiple trajectories, the log of 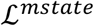 is calculated for each trajectory and summed over to obtain 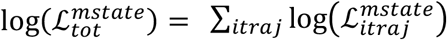.

### Position-dependent HMM

Thus far, the switching probabilities among different energy profiles have been considered to be independent of the motor position. However, chemical steps, such as catalytic reactions in motors, have been shown to depend on the position of motors^66^. Thus, it is desirable to make the switching probabilities position-dependent^67^. Consequently, we introduce a region called the switching region. Switching can only occur when the motor is in this region (Fig. 1 and Fig. 2A). The switching probability *p*(*s*_*i*-1_ → *s_i_*) in Eq. (1) can be replaced with the position-dependent switching probability *p*(*s*_*i*-1_ → *s_i_*|*x_i_*) to produce the corresponding likelihood 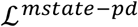 (“pd” stands for position-dependent). If *x_i_* is inside the switching region, it allows the state-switching transition. Otherwise, such transition is not allowed (bottom panel of Fig. 1B).

**Figure 2.**
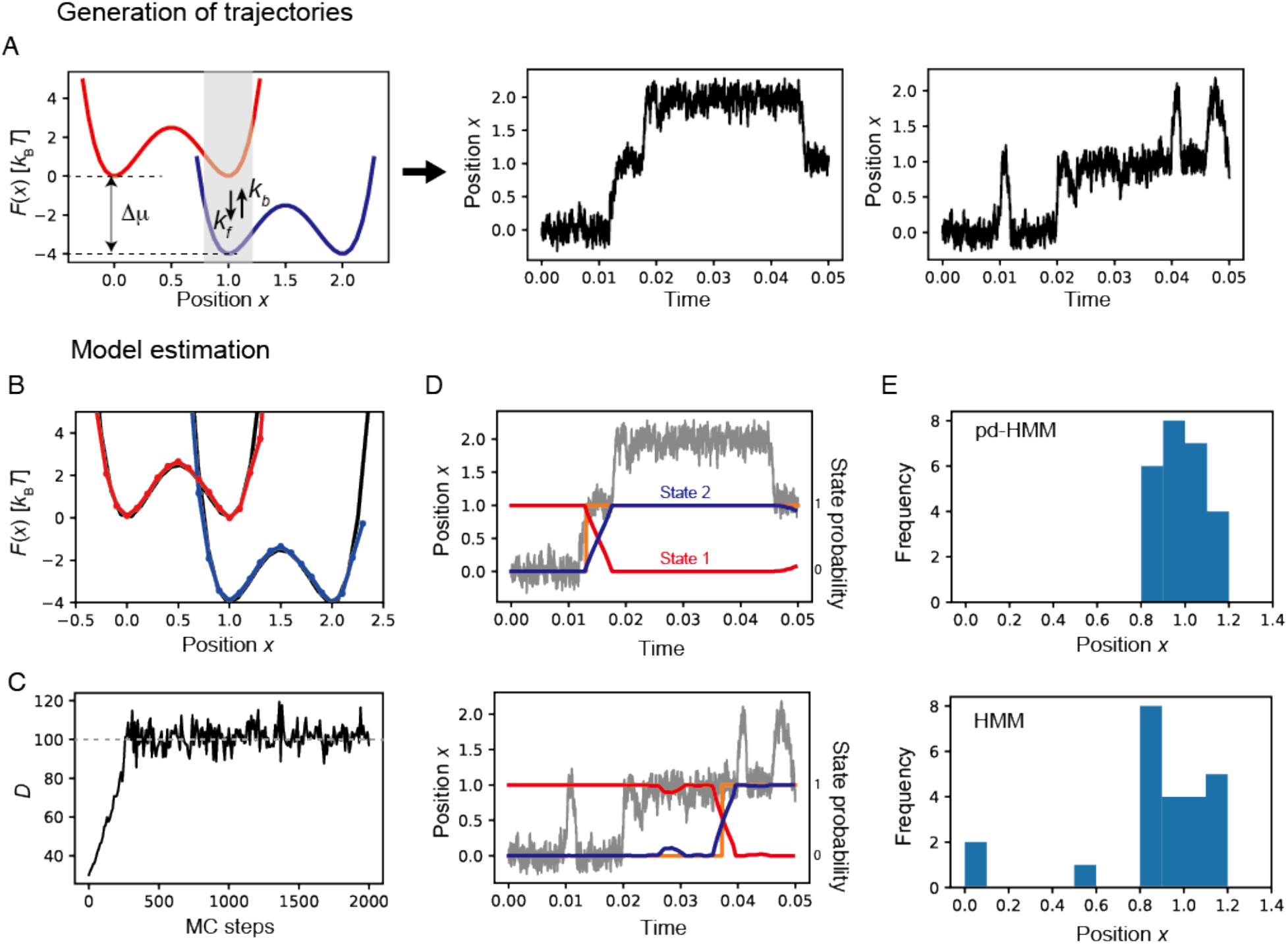
Generation of simulation trajectories and model estimation using the trajectories. (A) Two energy profiles used in Brownian dynamics simulation with stochastic switching (left panel) and two trajectories generated by the simulation (center and right panels). Switching only occurs in the shaded region of the left panel. (B) Estimated energy profiles shown as red and blue lines and true energy profile shown as black line. (C) Estimated diffusion coefficient during MC optimization steps (black line) and true diffusion coefficient (gray broken line). (D) pd-HMM predictions of state probabilities shown as red and blue lines and trajectories shown as gray lines. Orange lines represent actual timing of potential switching, where value 0 (1) corresponds to red (blue) potential. (E) Histograms of switching position obtained using pd-HMM (top panel) and HMM (bottom panel). Length and time units are arbitrary and can be interpreted as nanometers and seconds, respectively, from comparison with experiments.

From the HMM perspective, we effectively consider two different Markov models (or state-transition probabilities) depending on whether the motor is within or outside the switching region (bottom panel of Fig. 1B). This concept is implemented in the HMM forward-backward algorithm (see SI text). In the forward-backward algorithm, the likelihood and the state probability are calculated recurrently with one step in time in either the forward and backward direction. In this step, we introduce the position-dependent switching probabilities, as shown in the bottom panel of Fig. 1B. In this way, we obtain the likelihood 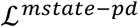 and the state probability that reflects the position-dependent switching probabilities. This model is termed position-dependent HMM (pd- HMM).

### Positional transition probability for diffusion model with a single energy profile

In this section, we describe the components needed to compute the likelihood function 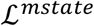. We relate the free energy profile and the diffusion coefficient to the positional transition probability, which was originally developed by Hummer and coworkers with extensive applications to simulation trajectories of protein folding^60,62,64^. We use a 1D model to calculate the transition probability between discretized positions (bins) from the underlying energy profile and the diffusion coefficient. The 1D Smoluchowski equation can be spatially discretized to yield the transition rate constant from bin *j* to bin *j* ± 1^68^:

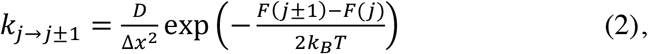

where *F*(*j*) is the free energy of bin *j, D* is the diffusion coefficient, and *Δx* is the bin size. We use this expression to construct a kinetic model in terms of the free energy and the diffusion coefficient. Note that for simplicity we do not consider the diffusion coefficient to be position-dependent in this study. The master equation in matrix form is 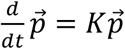, where *K* is the rate matrix, and 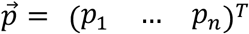 is the state probability vector for a total number of bins *n*. The rate matrix *K* is constructed from the transition rate constants as

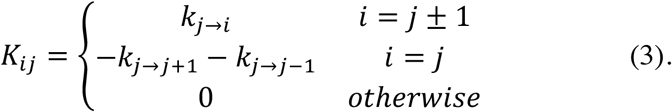

Then, the master equation can be formally solved to yield

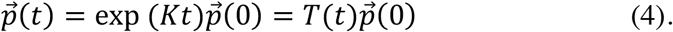

Each element of the transition matrix *T*(*t*) = exp (*Kt*) represents the positional transition probability, (*T*(*t*))_*ij*_ = *p*(*i, t|j*, 0), where *p*(*i, t|j*, 0) is the conditional probability of being in bin *i* at time *t*, given a previous location in bin *j* at time 0. Note that the subscript specifying the hidden state in *F* and *p*(*i,t*|*j*,0) is omitted here, because we consider only one state (that is, a single energy profile).

The eigenvalues of the rate matrix *K* are related to the timescales of the diffusion model^60^. The first non-trivial eigenvalue *λ*_2_ represents the slowest timescale of the model, where *λ*_1_ = 0 > *λ*_2_ > λ_3_. The implied timescale is expressed as *τ_i_* = – 1/*λ_i_*, where *i* ≥ 2.

### Translational symmetry and prior probability of model

As biomolecular motors have a catalytic cycle, the motor energy profiles are expected to possess translational symmetry with respect to the distance moved (or step size) *l_step_* after one cycle. Thus, the energy profile of a state is copied and shifted by the step size to represent the energy profile of the next state. The diffusion coefficient of the first state is also copied to the next state. With *n_shift_* = int(*l_step_*/Δ*x*) and *n* total number of bins, *F*_1_(0), …, *F*_1_(*n* – *n_shift_* – 1) are copied to *F*_2_(*n_shift_*), …, *F*_2_(*n* – 1) to ensure translational symmetry. Peripheral regions are defined as the parts of the energy profiles that are outside the lower/upper limit of the coordinate after shifting, that is, *F*_1_(*j*) with *j* = *n* – *n_shift_*, …, *n* – 1 and *F*_2_(*j*) with *j* = 0, …,*n_shift_* – 1. The peripheral regions, denoted as *j_s_* ∈ *out* with the subscript *s* representing the state, are expected to be rarely visited and have high energies. Thus, we use the prior probability that the energy of the peripheral regions increases at a constant rate:

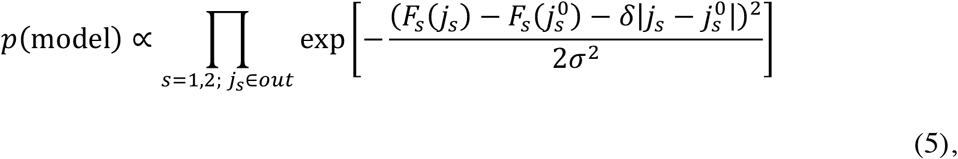

where *δ* is the increase in the energy per bin, 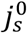 is the last bin before entering the region with 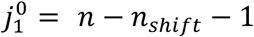 and 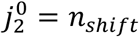, and *σ* is standard deviation that determines the strength of the restraint. Then, according to Bayes’ theorem, the posterior probability *p*(model\data) is

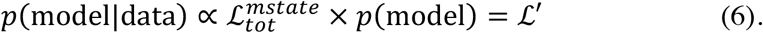

The posterior probability 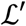 is maximized during the optimization process (see below).

### Maximizing posterior probability

Our objective is to optimize *F*(*x*) and *D* as well as the switching probabilities to maximize the posterior probability 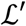 with respect to the input trajectory data. For this objective, Monte Carlo (MC) simulations were performed in the parameter space. Explicitly, the model parameters are *θ* = (*F*_1_(0),…, *F*_1_(*n* – 1),*F*_2_(0),…, *F*_2_(*n* – 1),ln *D*_1_, ln*D*_2_, *k_f_,k_b_*). The parameters *k_f_* and *k_b_* are related to the switching probabilities as *p*(1 → 2) = 1 – exp(–*k_f_* Δ*t*) and *p*(2 → 1) = 1 – exp(–*k_b_*Δ*t*), where Δ*t* is the lag time. In one MC step, each of these parameters is randomly changed as a trial move, 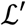 of the trial move is computed, and the trial move is accepted or rejected in comparison to the current 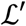. To ensure translational symmetry as described above, some parameters are copied between states. The widths of the trial moves are adjusted on the fly during the first half of the MC steps to obtain an acceptance ratio of ~0.3, which is then fixed during the second half. To compute the likelihood 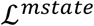, the positional transition probabilities are calculated for each state with given *F* and *D*, as described above, and then fed into HMM, as described above and Fig. 1B. The Metropolis criteria is used to accept or reject the trial move, 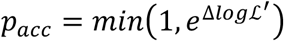, where 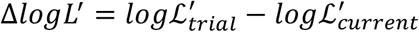^49,62^. In this way, the model parameters are guaranteed to be sampled based on the posterior probability 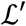.

### Generation of simulation trajectories

Based on Brownian dynamics, simulation trajectories for test analysis were generated numerically by the forward-Euler integrator,

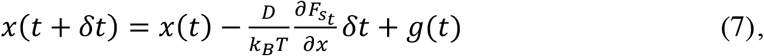

where *F_s_t__* is the potential energy of state *s_t_, D* is the diffusion coefficient, *δt* = 10^-6^ is the integration time step, and *g*(*t*) are Gaussian random numbers with mean zero and standard deviation 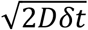. The model potential is *F_s_t__*(*x*) = *a*(*x* – (*s_t_* — 1))^2^(*x* – *s_t_*)^2^, where *s_t_* = 1 or 2 and *a* is a parameter controlling the barrier height. The state *s_t_*, and thus *F_s_t__*, is switched by the kinetic MC method with specified rate constants^69^ in a position-dependent manner as follows. The forward (*s_t_* = 1 → 2) and backward (*s_t_* = 2 → 1) switching events only occur within the range of 0.8 ≤ *x* ≤ 1.2 (Fig. 2A). Switching rates of *k_f_* = 200 and *k_b_* = 1 were used.

### RMSE of energy profile

The root-mean-squared error (RMSE) of an energy profile *Y*(*x*) relative to the true profile *Y*_0_ (*x*) is defined as

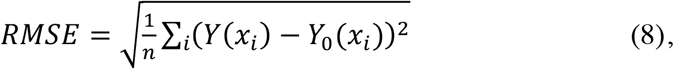

where *n* is the number of bins, excluding the peripheral regions. We shifted the entire profile *Y*(*x_i_*) by a constant *C* by the least squares method before calculating the RMSE, 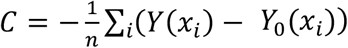.

### Alignment of chitinase trajectory segments

Each single-molecule trajectory of chitinase was divided into trajectory segments of two forward steps. The positional histogram of the segments was used to identify a high-frequency position for aligning the trajectory segments. A bin size of 0.15 nm was used to calculate the positional histogram for each trajectory segment. The high-frequency position was defined as a bin that was more frequent than the average frequency and the neighboring bins. The origin of the trajectory segments was set by subtracting the minimum of the identified high-frequency positions from the entire trajectory segment.

### Diffusion coefficient of gold nanoparticles

The diffusion coefficient of a 40-nm gold nanoparticle was estimated using the Stokes-Einstein relation,

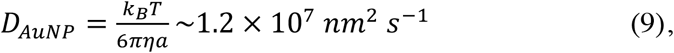

where *η* = 8.9 × 10^4^*kg m*^1^ *s*^1^ is the viscosity of water at 298 K, and *a* = 20 *nm* is the probe radius.

## RESULTS

### Reconstruction of diffusion model from simulation trajectories

We tested the developed method using simulation trajectories for which the true model parameters are known. Fig. 2A shows the two energy profiles used in the Brownian dynamics simulation with stochastic switching between these two profiles. The switching is position-dependent and only occurs in the shaded region (0.8 ≤ *x* ≤ 1.2). Two out of thirty independent trajectories generated by the simulation are also shown. The double-well potential is intended to mimic a scenario wherein chitinase moves back and forth before a catalytic reaction occurs^22^. Considering the size and time constant of chitinase stepping motions, the units of length and time in Fig. 2 can be interpreted as nanometers and seconds, respectively. The trajectories were generated by Brownian dynamics simulation for diffusive motions, and kinetic MC was used to switch between the two potentials with specified rate constants (see Methods). The generated trajectories were used to reconstruct the underlying energy profiles (*F*(*x*)) and the diffusion coefficient (*D*). In the estimation, the trajectories were discretized into bins with a bin size Δ *x* = 0.1 to produce 27 bins over the range −0.35 ≤ *x* ≤ 2.35, and the transitions among bins after a lag step of 400 were used as the observed data. In the Bayesian inference framework, the posterior probability 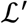 was maximized in terms of *F*(*x*) and *D* as well as the switching rates (see Methods). Note that the relative height of the two energy profiles (denoted as Δ*μ* in Fig. 2A) is arbitrarily shifted in the plots. Although the relative height Δ*μ* might be determined from the switching rates based on the detailed balance condition^70,71^, it is controversial that if nonequilibrium dynamics of biomolecular motors should satisfy the condition^32^.

The estimated energy profiles and diffusion coefficient are shown in Figs. 2B and 2C, respectively. In Fig. 2B, the estimated energy profiles averaged over the last 20% of the MC optimization steps are shown as red and blue lines, and the true profile is shown by a black line. The estimated diffusion coefficient along the MC optimization steps is shown in Fig. 2C, where the true value is shown as a broken line. Both the energy profiles and the diffusion coefficient agree quite well with the true profile (value). Furthermore, the state probabilities estimated using pd-HMM correctly reproduce the timing of the potential switching among the energy profiles (Fig. 2D). Considering that chemical steps, such as the hydrolysis reaction of chitin in chitinase, depend on the motor position, we look into position-dependency of the potential switching. The switching position was defined as the position at which the state probability of state 2 becomes larger than that of state 1, that is, *p_t_* (*s* = 2) > *p_t_*(*s* = 1), and its histogram was plotted in Fig. 2E. All the switching positions predicted by pd-HMM were inside the switching region (0.8 ≤ *x* ≤ 1.2), as one would expect (Fig. 2E, top). By contrast, the switching positions predicted by HMM without the position-dependent switching resulted in a few cases wherein the predicted switching position lay outside the switching region (Fig. 2E, bottom). This problem can be attributed to the relatively large lag step of 400, because the problem was almost fixed when a shorter lag step of 20 was used (Fig. S1). However, a short lag step resulted in an overestimation of the diffusion coefficient (as we discuss below). Moreover, the switching rates were reproduced by pd-HMM but underestimated by HMM (Fig. S2).

Then, we investigated how the accuracy of the estimated energy profiles and the diffusion coefficient depends on the lag time Δ*t* (or lag step) used to collect the positional transition data among bins. In Fig. 3A, the RMSE of the estimated profile (value) relative to the true profile (value) are plotted for different lag steps. We found that RMSE of the energy profile takes low values until lag step of 500 and then increases, whereas the RMSE of the diffusion coefficient converges at a lag step of approximately 400. Note that the diffusion coefficient was overestimated for short lag steps. This result suggests that it is necessary to use an appropriately long lag step, although an excessive length results in insufficient transition data. To determine the optimal lag time without knowledge of the true profiles, we performed a lag-time test to see how the implied timescale converges with the lag time^60,72^. The first non-trivial eigenvalue *λ*_2_ of the rate matrix estimated from the transition data with a prescribed lag time was converted into the so-called implied timescale using *τ*_2_ = −1/*λ*_2_ (see Methods). The implied timescale along the lag step converged at a lag step of approximately 500 (Fig. 3B). We only focus on the first non-trivial mode, because there is a clear timescale gap after this mode. Low RMSEs were obtained near the convergence point of the implied timescale (Fig. 3A and B). Thus, in the absence of knowledge of the true profiles, high accuracy can be achieved by using the lag time near the convergence point of the implied timescale.

**Figure 3.**
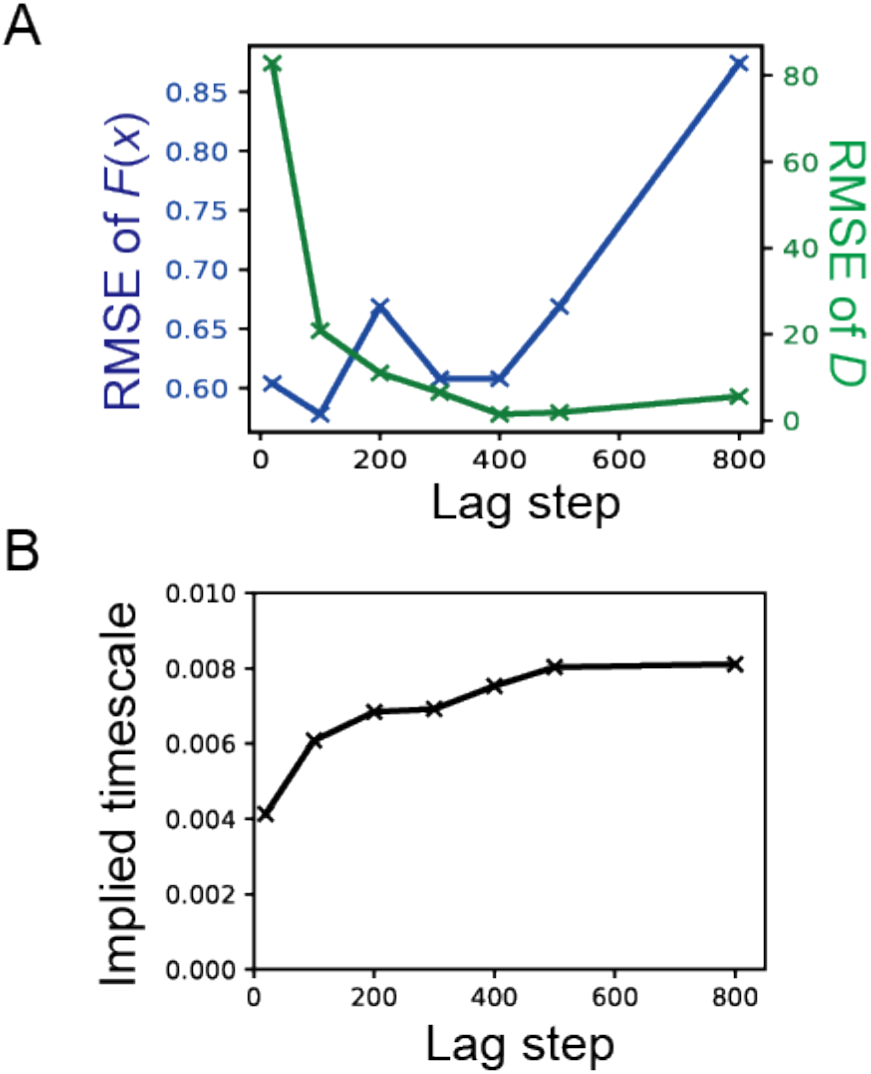
Effect of lag time on estimation accuracy. (A) Dependence of RMSEs of energy profile (*F*(*x*)) and diffusion coefficient (*D*) on lag step and (B) dependence of slowest implied timescale on lag step.

### Effect of background noise

In the previous section, we showed that our method accurately estimates the underlying energy profiles and diffusion coefficient from simulation trajectories. However, experimental singlemolecule trajectories always contain some background noise. We labeled chitinase with a 40-nm gold nanoparticle (AuNP) and observed 1-nm stepping motions using total internal reflection darkfield microscopy^22,73^: a localization precision of ~0.3 nm was obtained at a 0.333 – 0.5-ms time resolution. The localization precision is restricted by the detector saturation and can be improved^73^. However, the limited localization precision acts as background noise and should be addressed.

We added background noise to the simulation trajectories to investigate its effect on our predictions. Gaussian noise with zero mean and standard deviation of 0.3 nm was added to the *x* position saved every 500 steps (Δ*t* = 0.5 ms) to simulate the experimental background noise. A median filter with various window sizes was applied to reduce the noise level (Fig. 4A). The trajectories were then analyzed by our method. To address the noise, we used a larger bin size Δ*x* = 0.2 to produce a total of 13 bins over the range −0.3 ≤ *x* ≤ 2.3. Note that using this bin size in the analysis without noise accurately reproduces the energy profiles and diffusion coefficient (Fig. S3). It was prohibitively difficult to fully recover the true energy profiles and diffusion coefficient from the trajectories containing noise; however, partial recovery was achieved by adjusting the window size of the median filter. In the estimated energy profiles, the overall double-well shape with a lower energy barrier was recovered (Fig. 4B). Using trajectories median-filtered with a larger window size resulted in more accurate energy profiles. By contrast, increasing the window size produced an underestimated diffusion coefficient (Fig. 4C). Without applying the median filter, the diffusion coefficient was overestimated. Thus, this type of analysis can be used to determine lower and upper bounds on the diffusion coefficient. The timing of the potential switching was recovered reasonably well (Fig. 4D). Note that the standard deviation of 0.3 nm of the background noise is close to the step size of 1 nm, which makes an accurate estimation difficult. Irrespective of the limited estimation accuracy in the presence of background noise, we obtained the essential features of the energy profiles and a rough estimate of the diffusion coefficient.

**Figure 4.**
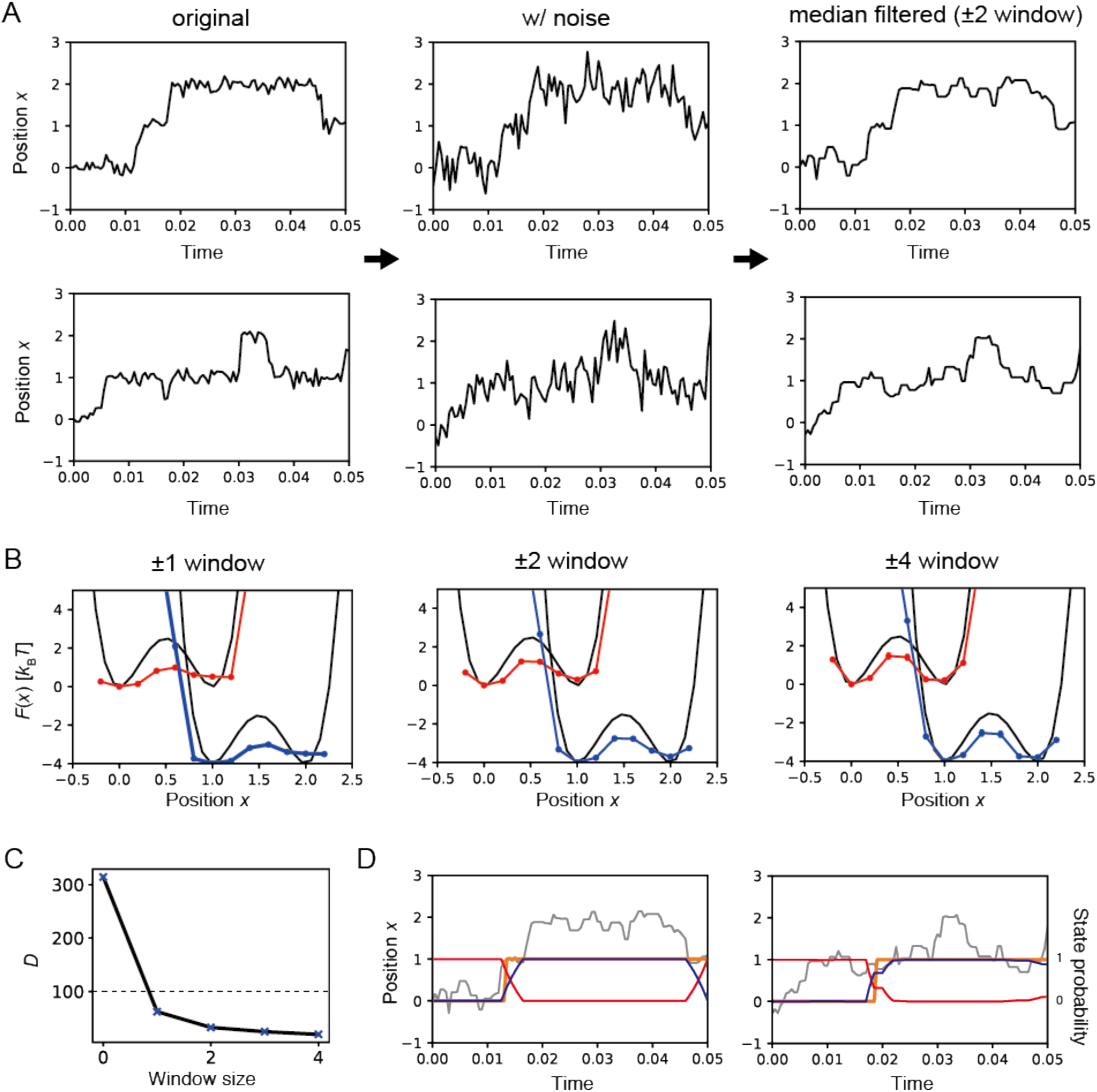
Effect of background noise. (A) Background noise was added to the original simulation trajectory and then reduced by a median filter. (B) Estimated energy profiles from trajectories median-filtered using window sizes of 1, 2 and 4, shown in left, center and right panels, respectively. (C) Estimated diffusion coefficients using trajectories median-filtered with different window sizes, where zero window size represents unfiltered trajectories, and dotted line represents the true value. (D) Two trajectories median-filtered with window size 2 are shown in gray lines.

Orange lines represent potential switching, where value 0 (1) corresponds to red (blue) potential. Estimated state probabilities of state 1 (red energy profile) and 2 (blue energy profile) shown by red and blue lines, respectively.

The prior probability appears to be an important factor in producing reliable estimates by the model from trajectories containing background noise. Using a weak prior probability (*σ* = 1 *k_B_ T, δ* = 1 *k_B_T*, see Methods) for trajectories containing background noise resulted in mixing of the two states, as evidenced by the state probabilities along the trajectories (Fig. S4). Consequently, the timing of switching was obscured for some trajectories. To rectify this situation, we used a stronger prior probability to analyze trajectories containing background noise (*σ* = 0.1 *k_B_T, δ* = 6 *k_B_T* for Fig. 4).

### Application to single-molecule trajectories of processive chitinase

We applied the developed method to single-molecule trajectories of processive chitinase. Processive chitinase is a unique biomolecular motor that moves unidirectionally on a crystalline chitin surface using the hydrolysis energy of a single chitin chain^74^. We previously used singlemolecule experiments to clarify the mechanochemical coupling scheme of processive chitinase (Fig. 5A) and proposed that it operates under a burnt-bridge Brownian ratchet mechanism, which is driven by a fast catalytic reaction and biased Brownian motions^22^. The burnt-bridge Brownian ratchet, has also been proposed for other motors, such as collagenase, that move processively on a collagen fibril with proteolysis energy^75,76^. Here, we aim to estimate the chemical-state-dependent free energy profile underlying the burnt-bridge Brownian ratchet mechanism of processive chitinase, directly from single-molecule trajectories.

**Figure 5.**
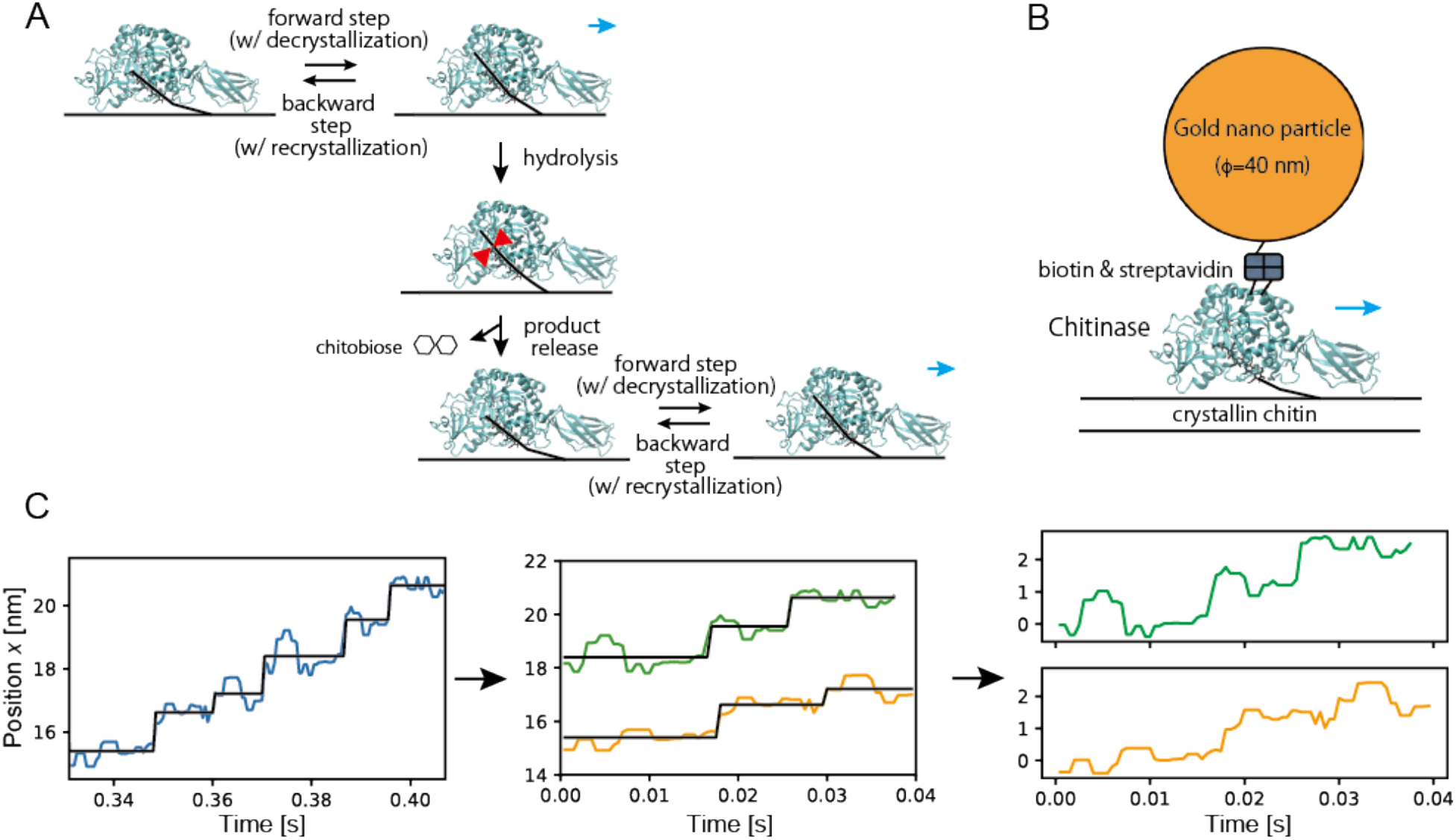
Single-molecule trajectories of processive chitinase. (A) Mechanochemical coupling scheme of chitinase A from bacteria *Serratia marcescens*, where cyan arrows represent direction of chitinase motion, and red saw teeth represent hydrolysis of a single chitin chain, shown as a black line. (B) Experimental setup of chitinase labeled with gold nanoparticle on crystalline chitin. (C) Single-molecule trajectory of chitinase at 2,000 frames per second (0.5-ms temporal resolution, left panel) was divided into two trajectory segments with two forward steps (center panel). Black lines represent steps identified by step-finding algorithm^77^. Trajectory segments were shifted to start from the origin (right panels)

In the single-molecule experiments, chitinase A from bacteria *Serratia marcescens* was labeled with a 40-nm gold nanoparticle (Fig. 5B), and total-internal reflection dark-field microscopy was used to observe the unidirectional motion of the particle on the chitin surface with a 0.3-nm localization precision and a 0.5-ms temporal resolution^22^. We used trajectories that were median-filtered to reduce positional noise. As the experimental trajectories were few and rather long in duration, the original trajectories were divided into segments to increase the sample size. In Fig. 5C, we show how the trajectory segments were generated. Each trajectory segment has two forward steps that were identified by the step-finding algorithm^22,77^. The segments were then aligned by matching the zero-points of the segments (see Methods).

The trajectory segments were analyzed in the same way as the test cases above. Since the trajectory segments have two forward steps, we assume two states (free energy profiles) that correspond to before and after the chemical state change (the hydrolysis reaction and the subsequent product release). This assumption is supported by molecular insights that each forward step corresponds to the sliding of the chitin chain through the chitinase cleft, after which the catalytic reaction and the subsequent product release reset the chitin-chitinase interaction (Fig. 5A). In the calculation, a bin size Δ *x* = 0.2 nm was used to produce 15 bins over the range of −0.5 ≤ *x* ≤ 2.5. To ensure translational symmetry, a shifting length of 1 nm was used, based on the chitinase step size (see Methods). The switching region 0.6 ≤ *x* ≤ 1.4 and the forward and backward switching rates of 100 s^-1^ and 0 s^-1^, respectively, were used. The switching region was based on the molecular insight that the hydrolysis reaction occurs at approximately *x* = 1. The forward switching rate was determined using the experimental time constants for the catalytic reaction (2.9 ms) and the subsequent product release (6.8 ms)^22^. The backward switching which is induced by the chitobiose binding and glycosidic bond formation at the catalytic site is unlikely to happen, because chitobiose concentration in solution is very low in our experimental condition. Therefore, the backward switching rate was set to zero.

The estimated diffusion model is shown in Fig. 6. The estimated free energy profiles capture the essential features of chitinase motions (Fig. 6A). Minima are located at approximately 0 and 0.6-1.0 nm for the pre-switching free energy profile and at approximately 1 and 1.6-2.0 nm for the post-switching free energy profile. These minima have similar free energies separated by a low barrier and are reflected in the chitinase stepping motions. After the chemical state change, the high-energy region *x* > 1.5 nm of the pre-switching profile decreases, resulting in a new minimum around the region of the post-switching free energy profile. The broad minimum after the barrier may explain variations in the step size. The estimated diffusion coefficient depends on the window size of the median filter (Fig. 6B). The larger the window size is, the smaller the estimated diffusion coefficient is. This result suggests a lower bound of ~200 nm^2^ s^-1^and an upper bound of ~1000 nm^2^ s^-1^ on the diffusion coefficient. Performing the same analysis on trajectories at a 0.333-ms temporal resolution increases the lower bound to ~500 nm^2^ s^-1^ (Fig. S5). These values are four orders of magnitude smaller than the calculated diffusion coefficient of the 40-nm gold probe *D_AuNP_* (see Methods), and explain the experimental observation that the chitinase velocity is not affected by the probe size^22^. The estimated time series of the state probabilities provide further information on the predictive ability of the model for switching between the two free energy profiles (Fig. 6C). These results suggest that chitinase moves forward via diffusive motion over a barrier, which facilitates the catalytic reaction for the forward switching of the free energy profile. The free energy profiles without a large driving gradient provide further evidence of the burnt-bridge Brownian ratchet mechanism of chitinase.

**Figure 6.**
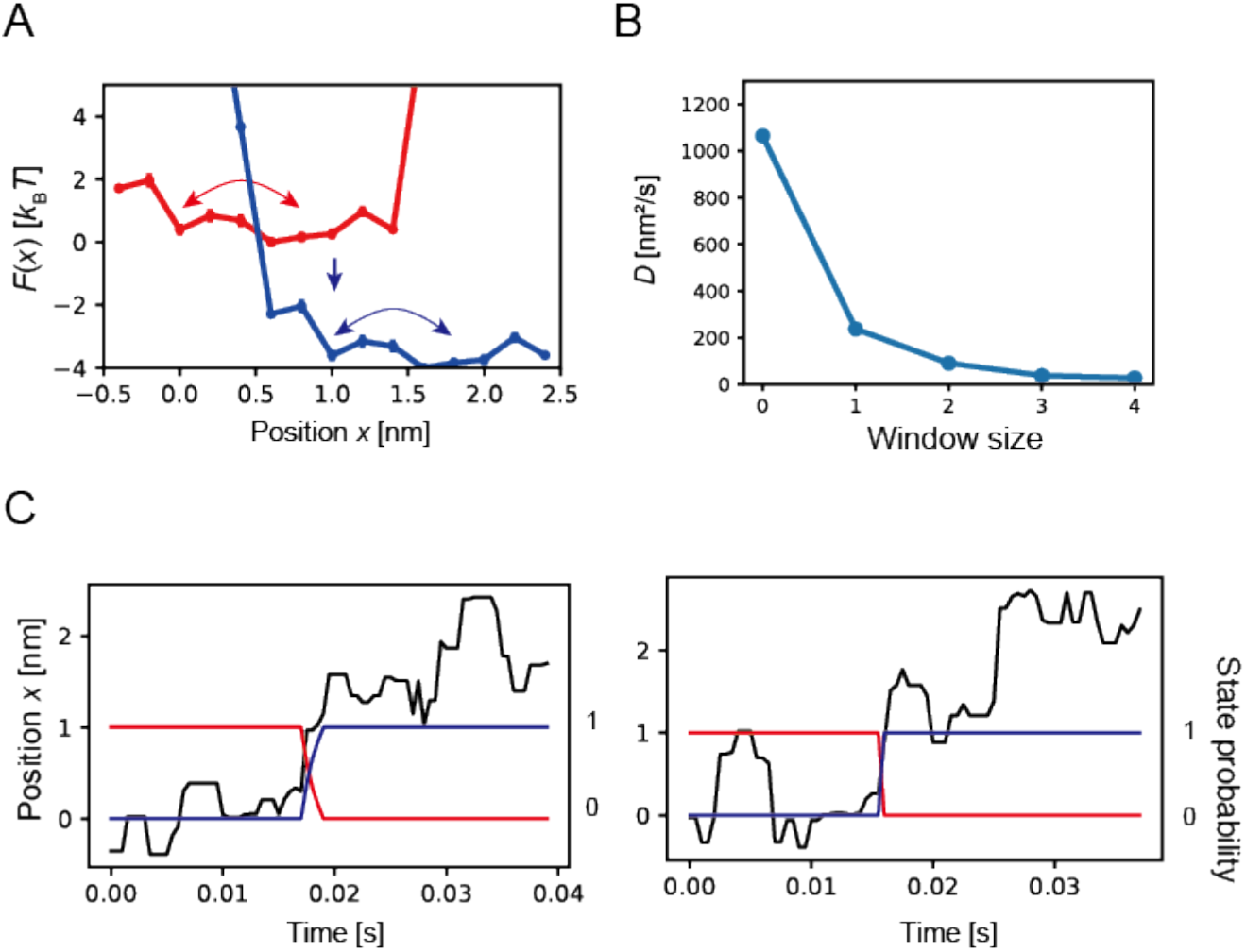
Estimated diffusion model of processive chitinase from trajectories at 0.5-ms temporal resolution. (A) Estimated free energy profiles before and after chemical state change are shown in red and blue, respectively, where arrows represent diffusive motion over a barrier and switching of energy profiles. (B) Estimated diffusion coefficients for different window sizes of median filter. (C) Examples of trajectory segments (black) and estimated state probability before (red) and after (blue) chemical state change are shown.

## DISCUSSION

The accuracy of the estimated chemical-state-dependent free energy profile and diffusion coefficient depends on the localization precision and temporal resolution of the single-molecule trajectory data. We used single-molecule trajectories of processive chitinase that were observed with a 0.3-nm localization precision and a 0.333 – 0.5-ms temporal resolution^22^. We selected trajectory segments showing clear stepping motions for the estimation. The clear stepping motions were used to align the trajectory segments with two forward steps used in the estimation (Fig. 5C).

We smoothed background noise by using a median filter with various window sizes. The test analysis of simulation trajectories containing background noise shows that our method would underestimate the barrier height and diffusion coefficient (Fig. 4B and C). Indeed, the estimated free energy profiles have a low barrier (Fig. 6A, Fig. S5A), which is likely caused by the background noise. Measurements with a higher localization precision and temporal resolution could be used in the future to improve the quality of the predictions^73^.

The friction coefficient estimated using the Einstein-Smoluchowski relation *ξ* = *k_B_ T/D* provides an insight into the energy conversion mechanism of processive chitinase. A stepping speed 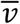 of processive chitinase can be estimated ~1000 nm s^-1^, considering that it takes ~1 ms (two frames with 0.5-ms temporal resolution) for the 1-nm stepping transitions^22^. Note that this is a rough estimate and for an accurate estimate one would need an improved localization precision and temporal resolution. Then, the viscous drag force 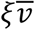 over a 1-nm stepping distance results in ~2 *k_U_T* of work. This work, compared to the hydrolysis energy of chitin (see below), leads to a low Stokes efficiency^78^, which is clearly different from the motion of the rotary motor F_1_-ATPase that shows high Stokes efficiency^79–82^. This result suggests that a considerable amount of the input hydrolysis energy of chitin is used for activities other than the mechanical movement of the motor. The important difference between the chitinase motor and typical ATPase motors is that the chitin rail shortens by one chitobiose unit during every catalytic cycle. Thus, it is essential to take the rail into consideration. Hypothetically considering the chitin rail alone, two reactions occur for every cycle: decrystallization of the chitobiose unit and the hydrolysis reaction (Fig. 5A). As decrystallization is an energy-consuming process, the net energy balance is maintained by using the input hydrolysis energy for the decrystallization of one chitobiose unit from crystalline β-chitin (the type of chitin used in the experiments). Although the free energies of hydrolysis of the β-1,4- glycosidic bond in chitin and decrystallization per chitobiose for β-chitin are not available to our best knowledge, the approximate free energies for similar reactions are available: the free energy of hydrolysis of the β-1,4-glycosidic bond in cellulose is −16 kJ mol^-1^ ≈ —6.5 *k_U_T*^83^, and the free energy of decrystallization per chitobiose for a-chitin is 5 kcal mol^-1^≈ 8 *k_U_T*^84^. The net energy balance of the entire chitinase-chitin system can be understood in a stepwise manner. In the catalytic cycle, starting from the state before decrystallization, processive chitinase decrystallizes the chitin chain and makes a forward step, where the chitin chain slides and attaches to the product binding site of chitinase. This binding free energy compensates the decrystallization energy, resulting in an equally stable state after the forward step^22^. Then, the hydrolysis reaction of the glycosidic bond and chitobiose release reset the system to the initial state, wherein the hydrolysis energy is liberated and used to expel the bound product. In summary, processive chitinase develops a stabilization mechanism for the high-energy intermediate formed in decrystallization, which is subsequently reset by the hydrolysis reaction.

Lastly, we mention previous methods used to estimate free energy profile(s) from nonequilibrium trajectories of the motors. Methods have been developed to estimate an effective single energy profile^31,85^. Although the dynamics generated by an effective single energy profile have been shown to reproduce some quantities, such as the mean number of steps per unit time^85^, these methods have limited applicability because chemical state changes that are central to the mechanochemical coupling of motors are not considered^3,86^. Toyabe and colleagues developed a pioneering method to estimate chemical-state-dependent free energy profile^87^. Although the method developed by Toyabe *et al*. is conceptually similar to our method, the likelihood function, the positional transition probability and the optimization process differed from those used in our method. In particular, the likelihood function (the path probability) used by Toyabe *et al*. only considers the best hidden-state path (as obtained by the Viterbi algorithm). Considering a computational cost of ~*O*(*N*) in both HMM and the Viterbi algorithm, the likelihood considering all the hidden-state paths used in our method should be more reliable. Another advantage offered by our method is that the free energy profiles and diffusion coefficient are calculated simultaneously, whereas the method of Toyabe *et al*. only optimizes free energy profiles with a fixed diffusion coefficient, which is estimated separately. In addition, our method can explicitly consider position-dependent switching rates.

## CONCLUSION

A chemical-state-dependent free energy profile can explain how biomolecular motors generate unidirectional motions. In this study, we introduced a framework to estimate the chemical-statedependent free energy profile and the diffusion coefficient from single-molecule trajectories of biomolecular motors. The method was tested using simulated trajectories and successfully reproduced the energy profiles and the diffusion coefficient. The method was then applied to single-molecule trajectories of processive chitinase. The estimated chemical-state-dependent free energy profile and diffusion coefficient provide a physical basis for the previously proposed burnt-bridge Brownian ratchet mechanism. The double-well free energy profile is shifted forward by the chemical state change to drive the unidirectional motion of chitinase (Fig. 6A). This mechanism is clearly different from the power-stroke mechanism, where a large driving gradient in the free energy profile is assumed. The estimated diffusion coefficient provides an insight into the energy conversion mechanism of processive chitinase.

## Supporting information

This file contains SI text and Figure S1-S5.

## ASSOCIATED CONTENT

### Supporting Information

The following files are available free of charge. The SI text describes the HMM algorithm. Figures S1-S5 show the model predictions.

## AUTHOR INFORMATION

### Author Contributions

K.O., A.N. and R.I. conceived the study. K.O. designed the algorithm and performed the research. K.O., A.N. and R.I. discussed the results. K.O. wrote the manuscript with contributions from A.N. and R.I.

## ACKNOWLEDGMENT

K.O. would like to thank Dr. Yohei Nakayama for discussions and Prof. Shoji Takada for comments on the manuscript. This study was supported by the Interdisciplinary Research Promotion Project (J281002 to K.O.) of the National Institutes of Natural Sciences (NINS), JSPS KAKENHI (JP18H02418 to R.I. and JP18H02415 to K.O.), and by Grant-in-Aid for Scientific Research on Innovative Areas “Molecular Engine” (grant number JP18H05424 to R.I.). K.O. was supported by the Building of Consortia for the Development of Human Resources in Science and Technology, MEXT, Japan.

## ABBREVIATIONS

HMM: hidden Markov model;
pd-HMM: position-dependent hidden Markov model;
1D model: one-dimensional model;
MC: Monte Carlo;
RMSE: root-mean-squared error

## TOC IMAGE

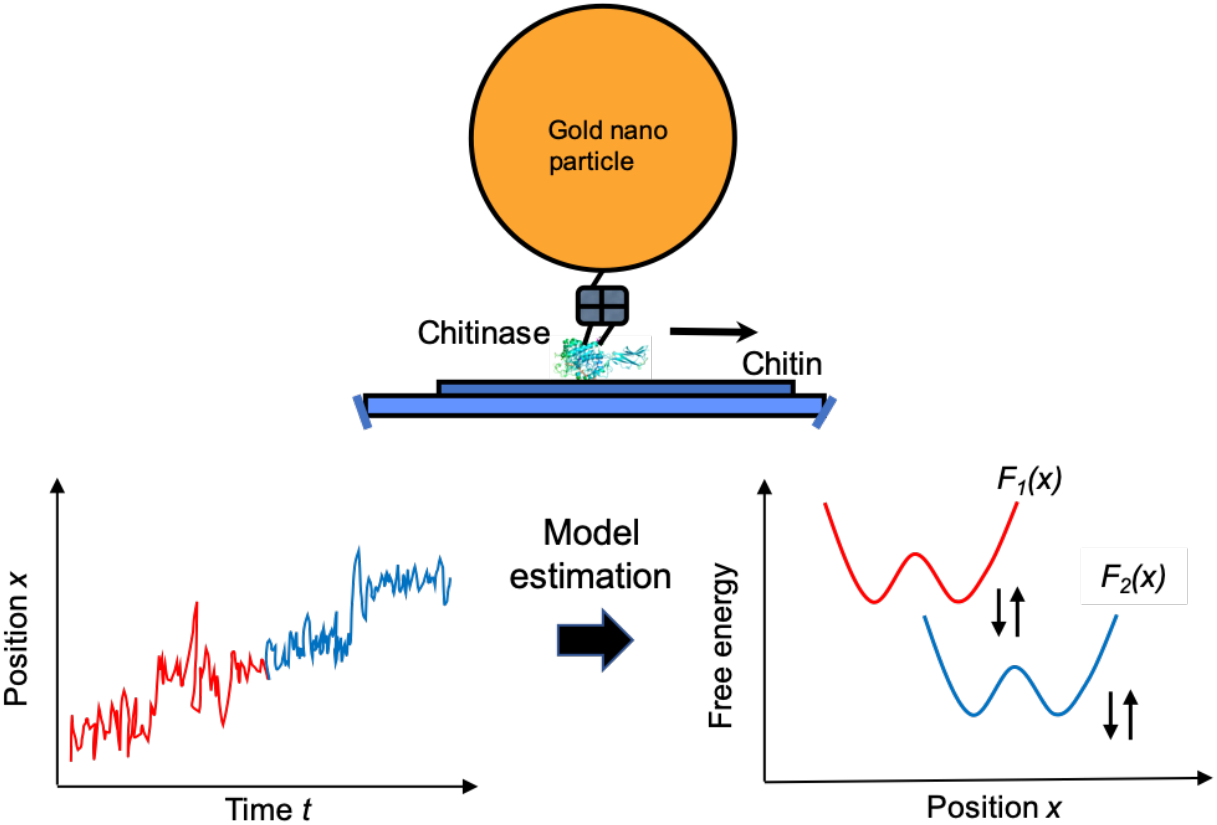

## Notes

#### Summary of Updates

Position-dependent HMM methods and results are included.

