## Supplementary material for "Chemical-state-dependent free energy profile from single-molecule trajectories of biomolecular motor: Application to processive chitinase": This file contains SI text and Figure S1-S5.

**SI text**

### The forward-backward algorithm of HMM

The likelihood function $\mathcal{L}^{mstate}$ introduced in the main text can be rewritten as,

$$\mathcal{L}^{mstate} = \sum_{S} \left[ p_{s_0}(y_0) \prod_{i=1}^{N-1} p(s_{i-1} \to s_i) \cdot p_{s_i}(y_i) \right]$$

where $y_i$ represents observed positional transition $x_i \to x_{i+1}$. Because possible number of hidden-state paths $S = (s_0, s_1, \cdots, s_{N-1})$ grows exponentially with number of time steps $N$ (that is, $2^N$ for the two state), direct calculation of the likelihood is computationally restrictive. In the forward-backward algorithm, the likelihood and state probability are calculated in a recursive manner to overcome the difficulty. In the forward direction, let $\alpha_t(j)$ denote probability of the observed data up to time *t*, given that it is in state *j* at time *t*. Thus,

$$\alpha_0(j) = p_j(y_0)$$

$$\alpha_t(j) = \sum_{s_0, \cdots, s_{t-1}} p_{s_0}(y_0) \cdot p(s_0 \to s_1) \cdot p_{s_1}(y_1) \cdots p(s_{t-1} \to j) \cdot p_j(y_t)$$

Then, $\alpha_t(j)$ satisfies the following recurrence relation,

$$\alpha_{t+1}(k) = \sum_{j} \alpha_t(j) \cdot p(j \to k) \cdot p_k(y_{t+1})$$

Note that this relation advances the likelihood calculation one step in time. After iteration over time steps $N$, the likelihood is obtained as $\mathcal{L}^{mstate} = \sum_j \alpha_{N-1}(j)$. Thus, the computational cost scales ~*O*(*N*).

In order to obtain the state probability, we consider similar recurrence relation in the backward direction. Let $\beta_t(j)$ denote probability of the observed data *after* time *t*, given that it is in state *j* at time *t*,

$$\beta_{N-1}(j) = 1$$

$$\beta_t(j) = \sum_{s_{t+1}, \cdots, s_{N-1}} p(j \to s_{t+1}) \cdot p_{s_{t+1}}(y_{t+1}) \cdots p(s_{N-2} \to s_{N-1}) \cdot p_{s_{N-1}}(y_{N-1})$$

Then, $\beta_t(j)$ satisfies the following recurrence relation,

$$\beta_{t-1}(k) = \sum_{j} p(k \to j) \cdot p_j(y_t) \cdot \beta_t(j)$$

Now, the probability of being in state $j$ at time $t$ (that is, the state probability) can be calculated as,

$$p_t(j) = \frac{\alpha_t(j)\beta_t(j)}{\sum_j \alpha_t(j)\beta_t(j)} = \frac{\alpha_t(j)\beta_t(j)}{L^{mstate}}$$

**Implementation of position-dependent HMM**

Here we consider the position-dependent switching. The likelihood function is,

$$\mathcal{L}^{mstate-pd} = \sum_S \left[ p_{s_0}(y_0) \prod_{i=1}^{N-1} p(s_{i-1} \to s_i | y_{i-1}) \cdot p_{s_i}(y_i) \right]$$

where $p(s_{i-1} \to s_i | y_{i-1})$ is the position-dependent switching probability. Note that $y_{i-1}$ represents the positional transition $x_{i-1} \to x_i$ and the switching probability only depends on $x_i$.

As we see above, the forward-backward algorithm advances one step in time in either the forward or backward direction. Thus, it is feasible to change the state-transition (or switching) probabilities based on the position at every iteration. As described in the main text, we only allow the state-switching transitions when the position is inside the switching region (pd-HMM in Fig. 1B). In this case, the recurrence relation in the forward direction is modified as,

$$\alpha_{t+1}(k) = \sum_j \alpha_t(j) \cdot p(j \to k \mid y_t) \cdot p_k(y_{t+1})$$

When $x_{t+1}$ (that is, the latter position of positional transition $y_t$) is inside the switching region (SR), which is denoted as $y_t \in SR$,

$$p(1 \to 1 \mid y_t \in SR) = e^{-k_f \Delta t}, \qquad p(1 \to 2 \mid y_t \in SR) = 1 - e^{-k_f \Delta t}$$

$$p(2 \to 1 \mid y_t \in SR) = 1 - e^{-k_b \Delta t}, \qquad p(2 \to 2 \mid y_t \in SR) = e^{-k_b \Delta t}$$

where $k_f$ and $k_b$ are switching rates for $1 \to 2$ and $2 \to 1$, respectively, and $\Delta t$ is lag time. Otherwise,

$$p(1 \to 1 \mid y_t \notin SR) = 1, \qquad p(1 \to 2 \mid y_t \notin SR) = 0$$

$$p(2 \to 1 \mid y_t \notin SR) = 0, \qquad p(2 \to 2 \mid y_t \notin SR) = 1$$

In summary, the state-changing transition at time between $y_t$ and $y_{t+1}$ observations is allowed only if the latter position of $y_t$ is inside the SR. We consider the same position-dependent switching probability for the backward direction. Then, the likelihood and state probability obtained by the forward-backward algorithm reflects the position-dependent switching rates.

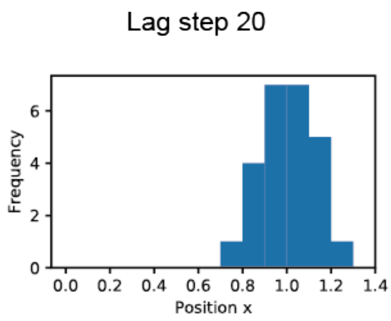


**Figure S1.** Histogram of the switching position by HMM with lag step 20.

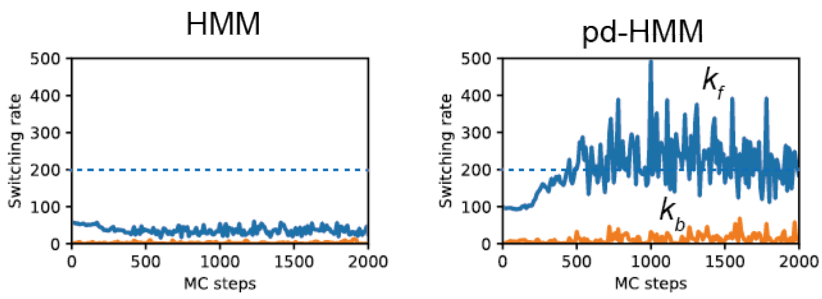


**Figure S2.** The switching rates estimated by HMM (left) and by pd-HMM (right) during the MC optimization steps. The blue dotted lines represent the true value of $k_f$.

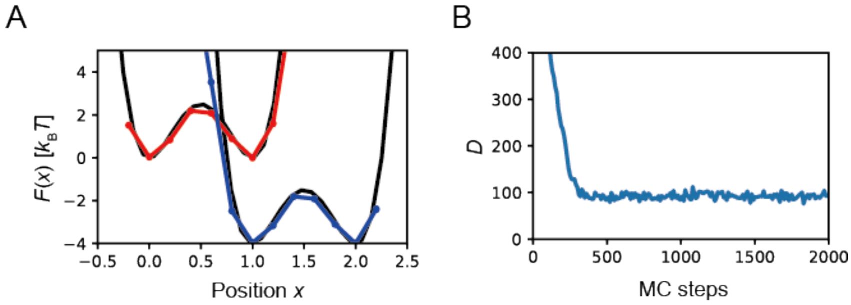


**Figure S3.** Estimated diffusion model with the bin size $\Delta x = 0.2$. (A) Estimated energy profiles are shown. (B) Estimated diffusion coefficient along MC steps is shown.

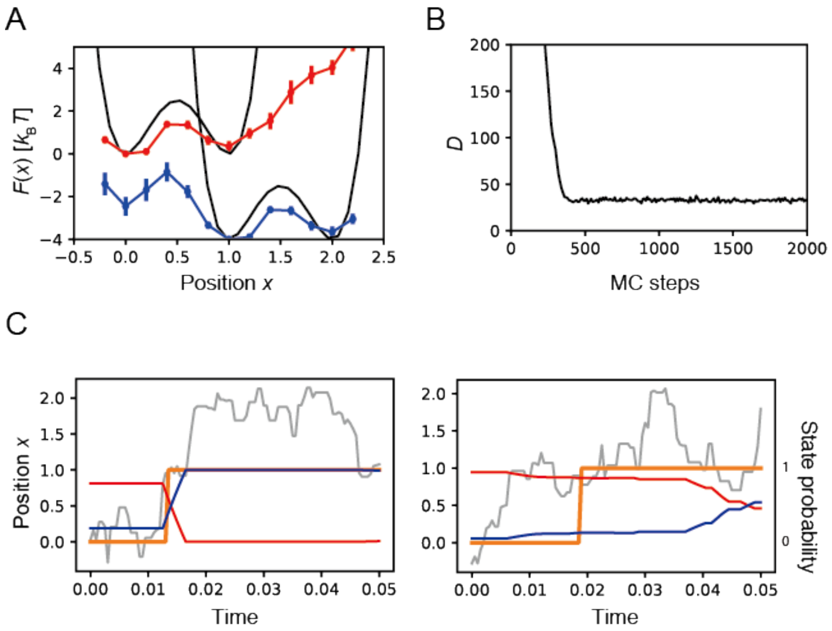


**Figure S4.** Estimated model with a weak prior probability ($\sigma = 1\ k_B T$ and $\delta = 1\ k_B T$) from trajectories with background noise median-filtered with window size 2. (A) Estimated energy profiles. (B) Estimated diffusion coefficient. (C) Estimated state probability along trajectories.

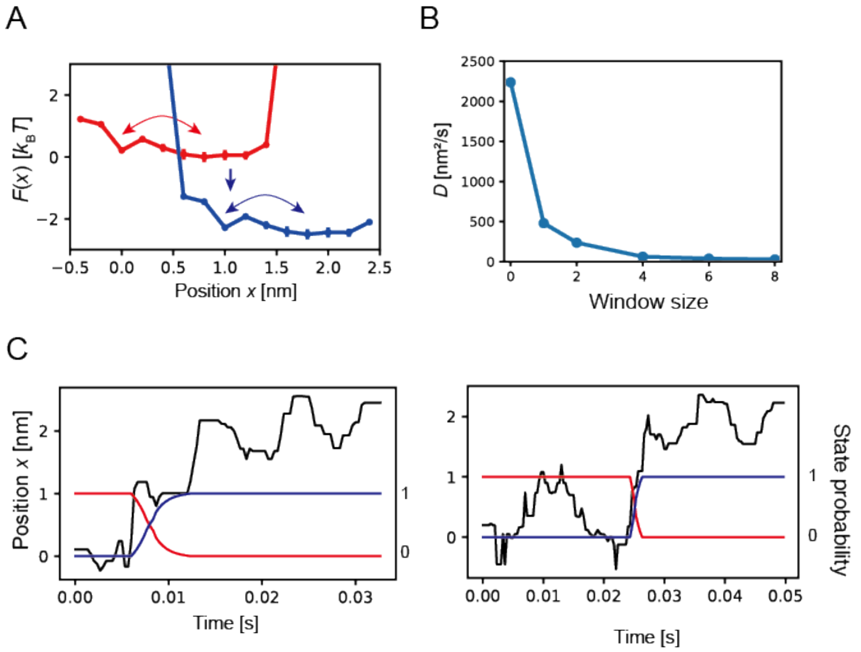


**Figure S5.** Estimated diffusion model of chitinase from trajectories at 0.333 ms temporal resolution. (A) Estimated free energy profiles of before and after the chemical-state change are shown in red and blue, respectively. Arrows represent diffusive motions over a barrier and switching of energy profiles. (B) Estimated diffusion coefficients with different window sizes of the median filter. (C) Examples of trajectory segments (black) and estimated state probability of before (red) and after (blue) the chemical state change are shown.
